# The kleisin subunit controls the function of meiotic cohesins by determining the mode of DNA binding and differential regulation by SCC-2 and WAPL-1

**DOI:** 10.1101/2022.10.12.511771

**Authors:** Maikel Castellano-Pozo, Georgios Sioutas, Consuelo Barroso, Pablo Lopez-Jimenez, Angel Luis Jaso-Tamame, Oliver Crawley, Nan Shao, Jesus Page, Enrique Martinez-Perez

## Abstract

The cohesin complex plays essential roles in chromosome segregation, 3D genome organisation, and DNA damage repair through its ability to modify DNA topology. In higher eukaryotes, meiotic chromosome function, and therefore fertility, requires cohesin complexes containing meiosis-specific kleisin subunits: REC8 and RAD21L in mammals and REC-8 and COH-3/4 in *C. elegans*. How these complexes perform the multiple functions of cohesin during meiosis and whether this involves different modes of DNA binding or dynamic association with chromosomes is poorly understood. Combining time-resolved methods of protein removal with live imaging and exploiting the temporospatial organisation of the *C. elegans* germline, we show that REC-8 complexes provide sister chromatid cohesion (SCC) and DNA repair, while COH-3/4 complexes control higher-order chromosome structure. High-abundance COH-3/4 complexes associate dynamically with individual chromatids in a manner dependent on cohesin loading (SCC-2) and removal (WAPL-1) factors. In contrast, low-abundance REC-8 complexes associate stably with chromosomes, tethering sister chromatids from S-phase until the meiotic divisions. Our results reveal that kleisin identity determines the function of meiotic cohesin by controlling the mode and regulation of cohesin-DNA association, and are consistent with a model in which SCC and DNA looping are performed by variant cohesin complexes that coexist on chromosomes.

## Introduction

Cohesin belongs to a family of structural maintenance of chromosomes (SMCs) protein complexes that are essential components of mitotic and meiotic chromosomes due to their ability to modify the topology of DNA (Yatskevich et al., 2019). In somatic cells, cohesin ensures correct chromosome segregation by providing sister chromatid cohesion (SCC) between S-phase and the onset of anaphase, ensures genome stability through its role in DNA damage repair, and regulates gene expression by controlling genome folding through DNA looping. In addition, during meiosis, cohesin orchestrates the formation of proteinaceous axial elements that promote pairing and recombination between homologous chromosomes, and facilitates the two-step release of SCC during the consecutive meiotic divisions. As inter-homologue recombination and the step-wise release of SCC are prerequisites for the formation of euploid gametes, the correct function of meiotic cohesin is essential for fertility.

The core of cohesin consists of a ring-shaped complex formed by two SMC proteins (Smc1 and Smc3) and a kleisin subunit that bridges the ATPase heads of Smc1 and Smc3. A family of HAWK (Heat repeat proteins Associated With Kleisin) proteins are recruited by the kleisin to control the loading, removal, and activity of cohesin on DNA (Wells et al., 2017). These include Scc3, Pds5, and the cohesin loader Scc2, which is required for the association of cohesin with mitotic and meiotic chromosomes (Ciosk et al., 2000; Lightfoot et al., 2011). Pds5 and Scc3 in turn interact with Wapl, a factor that promotes removal of cohesin throughout the cell cycle in a manner that allows reloading of removed complexes (Kueng et al., 2006). Cleavage of the kleisin subunit by separase at anaphase onset during the mitotic and meiotic divisions triggers irreversible cohesin removal and release of SCC to promote chromosome segregation (Buonomo et al., 2000; Uhlmann et al., 1999). Thus, the kleisin subunit plays a key role in controlling cohesin’s action on DNA.

An essential aspect of meiotic chromosome morphogenesis is the loading of cohesin complexes in which the mitotic kleisin Scc1 is replaced by the meiosis-specific kleisin Rec8 (Klein et al., 1999; Watanabe and Nurse, 1999). Substitution of Rec8 by Scc1 during yeast meiosis results in severe defects during meiotic prophase and in premature loss of SCC during the first meiotic division (Brar et al., 2009; Toth et al., 2000), evidencing that Rec8 cohesin is functionally different from Scc1 cohesin. Higher eukaryotes express additional meiosis-specific kleisins beyond REC8, including RAD21L in mammals and the highly identical and functionally redundant COH-3 and COH-4 in *C. elegans*, which appear to be the functional counterpart of mammalian RAD21L (Gutierrez-Caballero et al., 2011; Ishiguro et al., 2011; Lee and Hirano, 2011; Severson et al., 2009; Severson and Meyer, 2014). How complexes differing in their kleisin subunit perform the multiple roles of meiotic cohesin remains poorly understood. Moreover, the dynamic association of cohesin with chromosomes is key for some of cohesin’s functions in somatic cells (Tedeschi et al., 2013), but whether meiotic cohesins associate dynamically with chromosomes is not known. In this study, we exploit the experimental advantages of the *C. elegans* germline to investigate how REC-8 and COH-3/4 complexes contribute to different aspects of meiotic cohesin function and whether this involves dynamic association of these complexes with meiotic prophase chromosomes.

## Results and discussion

### Dynamically-bound COH-3/4 complexes are the main organisers of axial elements

We first set up to determine the relative abundance of the three meiotic kleisins (REC-8, COH-3, and COH-4) on the axial elements of three-dimensionally intact pachytene nuclei. The endogenous *rec-8, coh-3*, and *coh-4* loci were individually tagged with GFP by CRISPR to generate three strains homozygous for GFP-tagged versions of each of the meiotic kleisins. GFP tagging of meiotic kleisins did not affect protein functionality as strains homozygous for *rec-8::GFP*, or for both *coh-3::GFP* and *coh-4::GFP* showed normal chiasma formation (Figure 1A). To quantify the signal intensity of meiotic kleisins in pachytene nuclei, germlines were dissected, stained with anti-GFP antibodies, and imaged under the same exposure conditions. Half nucleus projections were made to prevent overlap between axial elements from different chromosomes, followed by quantification of the fluorescent signal. This demonstrated that COH-3 is the most abundant kleisin on pachytene axial elements, which contain similar amounts of COH-4 and REC-8 (Figure 1B). The ratio of COH-3/4 to REC-8 cohesin is 3.5, consistent with the observation that COH-3/4 cohesin plays a more prominent role than REC-8 cohesin in axis integrity of pachytene nuclei (Castellano-Pozo et al., 2020) and with previous estimations of REC-8 and COH-3/4 abundance on spread pachytene nuclei (Woglar et al., 2020). To explore the contribution of REC-8 and COH-3/4 cohesin to the organization of axial elements, we used structural illumination microscopy (SIM) to image SMC-1::GFP (a cohesin SMC subunit common to all types of cohesin in worms) in germlines of controls, *rec-8* single, and *coh-3 coh-4* double mutants. Continuous axial elements labelled by SMC-1::GFP were observed in wild-type controls and *rec-8* mutants, while SMC-1::GFP signals appeared as discontinuous weak signals in pachytene nuclei of *coh-3; coh-4* double mutants (Figure 1C). Thus, by ensuring the integrity of axial elements, COH-3/4 complexes are key regulators of higher-order chromosome organisation during meiotic prophase.

**Figure 1.**
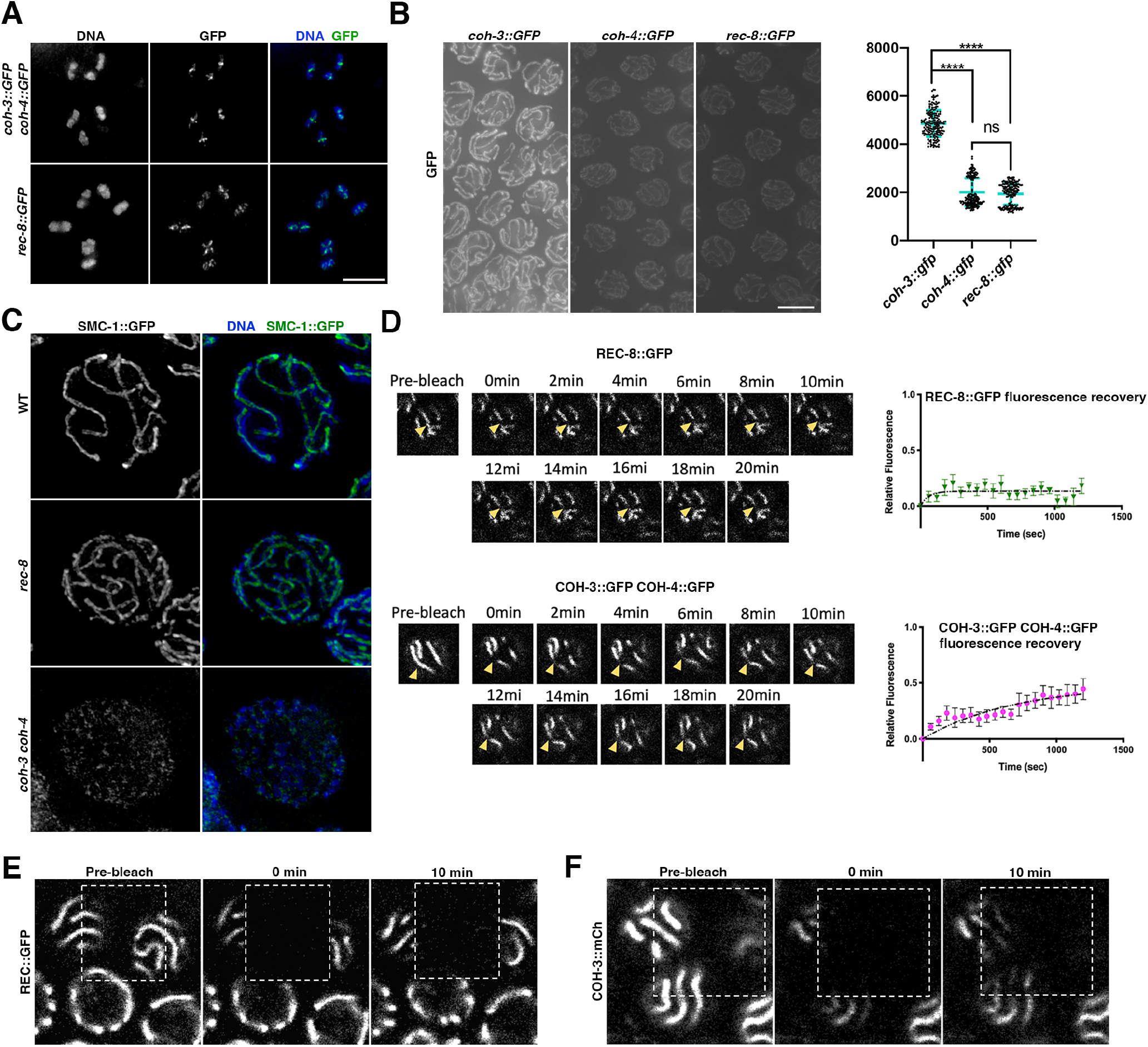
Highly abundant COH-3/4 complexes promote axis integrity and associate dynamically with pachytene chromosomes. **(A)** Projections of diakinesis oocytes stained with DAPI and anti-GFP antibodies to visualise COH-3::GFP COH-4::GFP (top panel) or REC-8::GFP (bottom panel). The presence of 6 bivalents confirms normal chiasma formation. **(B)** Non-deconvolved projections of pachytene nuclei of indicated genotype stained with anti-GFP antibodies. Images were acquired under the same exposure conditions and panels were adjusted with the same settings to allow direct comparison of GFP intensity in the different genotypes. Graph shows anti-GFP intensity quantification, error bars indicate mean and SD, p values were calculated using a two-tailed Mann-Whitney U test. N (number of axes analysed (1 or 2 per nucleus): 192 (*coh-3::GFP*), 200 (*coh-4::GFP*), 191 (*rec-8::GFP*). **(C)** projections of pachytene nuclei from worms expressing SMC-1::GFP of indicated genotypes stained with anti-GFP antibodies and DAPI and imaged by SIM. Note the presence of continuous-linear structures containing SMC-1::GFP in WT and *rec-8* mutants, but not in *coh-3 coh-4* double mutants. **(D)** FRAP analysis of REC-8::GFP and COH-3::GFP COH-4::GFP in pachytene nuclei. Images show pre-and post-photobleaching images at indicated time points with arrowheads indicating the photobleached area on the axial element that was followed through the experiment. As nuclei move through the experiment, the focal plane was adjusted to follow the indicated region of the bleached axial element, while other regions can be out of focus. N= 9 nuclei (REC-8::GFP), n= 11 (COH-3::GFP COH-4::GFP). Error bars indicate SEM. **(E)** High resolution FRAP images of worms expressing REC-8::GFP (genotype: *rec-8::GFP rec-8Δ*) at indicated times before and after photobleaching the area indicated by the dashed rectangle. Note that 10 minutes after photobleaching there is no recovery of REC-8::GFP signal on bleached axial elements. **(F)** High resolution FRAP images of worms expressing COH-3::mCherry and REC-8::GFP (genotype: *coh-3::mCh coh-3Δ rec-8::GFP rec-8Δ*) at indicated times before and after photobleaching the area indicated by the dashed rectangle. Note that 10 minutes after photobleaching there is recovery of COH-3::mCh signal on bleached axial elements. Scale bar= 5 *μ*m in all panels.

We next addressed whether REC-8 or COH-3/4 complexes associate dynamically with meiotic prophase chromosomes by performing Fluorescence Recovery After Photobleaching (FRAP) in pachytene nuclei of live animals. Following photobleaching of a small region of an axial element, we observed very little recovery of axis-associated REC-8::GFP signal, evidencing little reloading of REC-8::GFP over the imaging period (20 minutes) (Figure 1D). In contrast, FRAP analysis of pachytene nuclei from both *coh-3::GFP* and *coh-3::GFP coh-4::GFP* homozygous worms demonstrated regaining of fluorescence signal on the photobleached region of the axis, which approached 50% of pre-bleach levels over 20 minutes (Figures 1D and 2A). To further confirm these observations, we also performed high-resolution FRAP experiments (see methods) in which we bleached a larger area spanning two nuclei and acquired images at only three times points (pre-bleach, 0 minutes, and 10 minutes post-bleach) using transgenic strains expressing fully functional fluorescently-tagged versions of REC-8 and COH-3 (Castellano-Pozo et al., 2020). Consistent with the full time-course experiment shown above, high-resolution FRAP demonstrated clear reloading of COH-3::mCherry, but not REC-8::GFP to axial elements of pachytene nuclei 10 minutes after photo bleaching (Figures 1E-F). Thus, in contrast with REC-8 cohesin, COH-3/4 complexes associate dynamically with pachytene axial elements. In mammalian somatic cells the creation of a stable pool of cohesin requires passage through S-phase (Gerlich et al., 2006) and in worms only REC8 complexes associate with chromosomes during meiotic S-phase, while COH-3/4 load post S-phase (Severson and Meyer, 2014). Therefore, the different loading time of REC-8 and COH-3/4 cohesin may be an important determinant in the dynamics of these complexes. As RAD21L complexes also associate post S-phase during mouse meiosis (Ishiguro et al., 2014), our results suggest that, similar to COH-3/4, RAD21L may also associate dynamically with pachytene chromosomes.

### WAPL-1 and SCC-2 promote the dynamic association of COH-3/4 cohesin with pachytene chromosomes

The dynamic association of COH-3/4 cohesin with pachytene axial elements led us to test whether factors that promote loading and removal of cohesin during the mitotic cell cycle also control COH-3/4 dynamics. In mammalian somatic cells, WAPL is required to maintain a pool of cohesin that associates dynamically with chromosomes (Tedeschi et al., 2013), and we have previously shown that in the absence of WAPL-1 the levels of axis-associated COH-3/4 are increased (Crawley et al., 2016), suggesting that WAPL-1 may control the dynamic association of COH-3/4 complexes. To determine if this was the case, we performed FRAP in *wapl-1* mutant worms. While the dynamics of REC-8 GFP were similar in the presence and absence of WAPL-1, showing almost no increase of post-photobleaching fluorescence intensity, in the case of COH-3::GFP and COH-4::GFP the increase in post-photobleaching intensity was largely lost in the absence of WAPL-1 (Figure 2A). Therefore, WAPL-1 is required to ensure the dynamic association of COH-3/4 cohesin with axial elements in pachytene nuclei. WAPL also induces cohesin removal during late meiotic prophase in yeast and plants, but whether this also occurs at earlier stages is not currently understood (Challa et al., 2019; De et al., 2014). Removal of WAPL in mouse oocytes at the end of meiotic prophase results in increased binding of SCC1 cohesin, but not REC8 cohesin, suggesting that REC8 cohesin is also refractory to WAPL-mediated removal during mammalian meiosis (Silva et al., 2020). Elucidating the mechanisms that protect REC8 complexes in mammals and worms from WAPL-mediated removal remains an important question.

**Figure 2.**
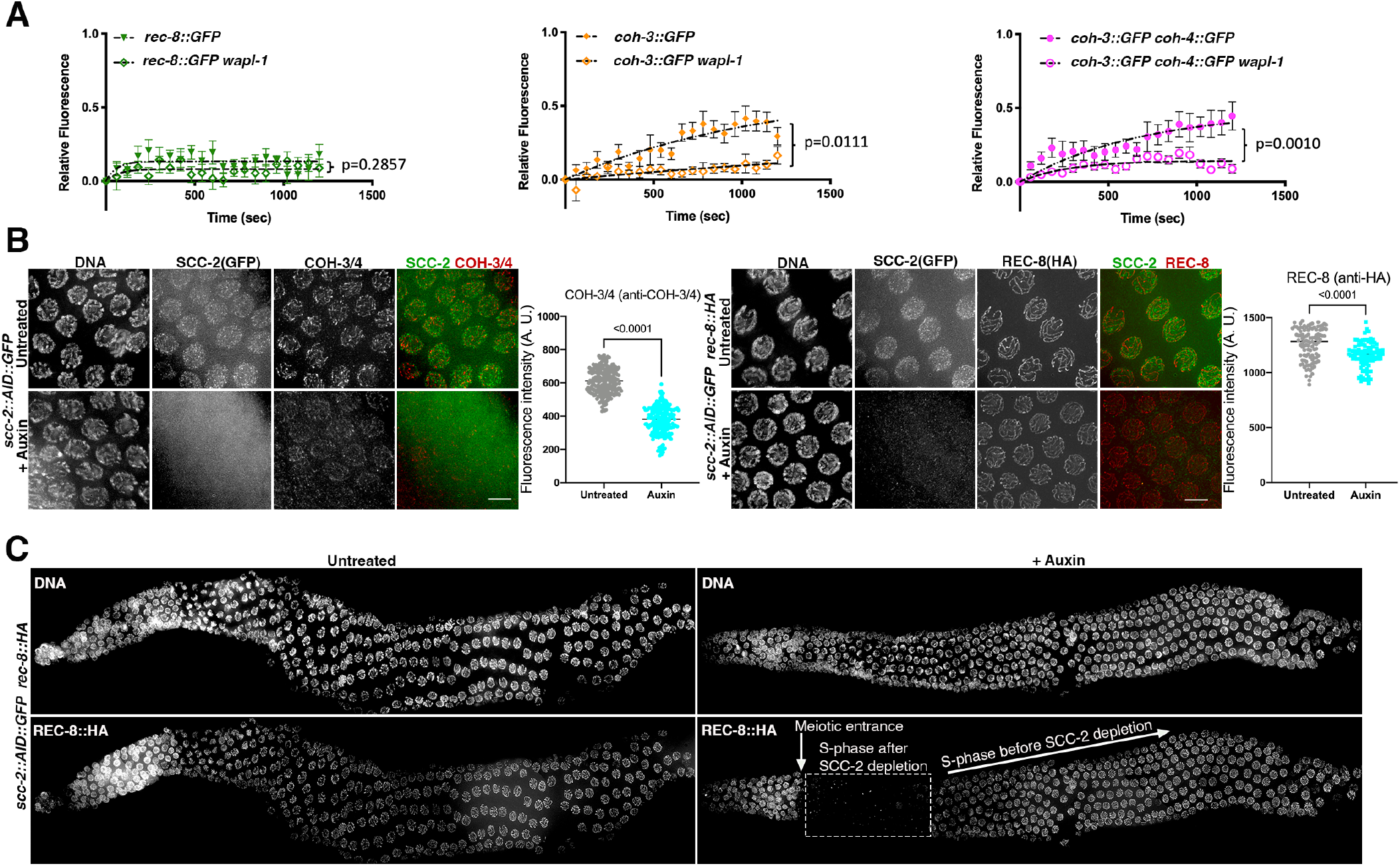
WAPL-1 and SCC-2 control the dynamic association of COH-3/4, but not REC-8, complexes with pachytene chromosomes. **(A)** FRAP analysis of REC-8::GFP, COH-3::GFP, and COH-3::GFP COH-4::GFP in pachytene nuclei of WT and *wapl-1* mutants. Data for WT REC-8::GFP and WT COH-3::GFP COH-4::GFP is the same as shown in Figure 1D. Number of nuclei analysed: n= 9 nuclei (REC-8::GFP) n= 10 (REC-8::GFP *wapl-1*), n=10 (COH-3::GFP) n= 10 (COH-3::GFP *wapl-1*), n= 11 (COH-3::GFP COH-4::GFP) n= 10(COH-3::GFP COH-4::GFP *wapl-1*). Error bars indicate SEM and p values were calculated by Mann-Whitney tests between the Ymax predicted values of the one-phase association curves of individual experiments. **(B)** Projections of pachytene nuclei from *scc-2::AID::GFP* (left) and *scc-2::AID::GFP rec-8::HA* (right) from control worms (untreated) and from worms treated with 4 mM auxin for 14 hours. SCC-2::GFP was visualised using anti-GFP antibodies, COH-3/4 using anti-COH-3/4 antibodies, and REC-8 using anti-HA antibodies. Note efficient removal of SCC-2::AID::GFP signal following auxin treatment, which induces strong decrease of COH-3/4 but not REC-8 signal. Graphs show quantification of overall nuclear intensity in projections of pachytene nuclei before and after auxin treatment. Number of nuclei analysed: COH-3/4 (225 untreated, 223 +auxin) REC-8 (105 untreated, 89 +auxin), lines indicate median and p values were calculated by two-tailed Mann Whitney U test. **(C)** Whole germline projections of *scc-2::AID::GFP rec-8::HA* worms stained with DAPI and anti-HA antibodies. Note the absence of REC-8-labelled axial elements in the region indicated by the dashed rectangle, corresponding to nuclei that entered meiosis after SCC-2 depletion. Nuclei that underwent meiotic S-phase before SCC-2 depletion retain strong REC-8::HA staining on axial elements. Scale bar= 5 *μ*m in all panels.

The fact that COH-3/4 cohesin complexes associate *de novo* with axial elements during pachytene led us to evaluate whether SCC-2, which is required for cohesin loading at the onset of meiosis and localizes to pachytene axial elements (Lightfoot et al., 2011), controls this process. As auxin-mediated protein degradation allows cohesin depletion from pachytene nuclei (Castellano-Pozo et al., 2020; Zhang et al., 2015), we tagged the endogenous *scc-2* locus with GFP and AID tags at its C-terminus using CRISPR. We reasoned that if SCC-2 promotes reloading of COH-3/4 complexes that are removed by WAPL-1, then, depletion of SCC-2 following normal cohesin loading during early meiosis should induce a decrease of COH-3/4 cohesin in pachytene nuclei. As nuclei take about 36 hours to progress from meiotic S-phase to late pachytene (Jaramillo-Lambert et al., 2007), treating *scc-2::AID::GFP* homozygous worms with auxin for 14 hours should result in germlines containing mid and late pachytene nuclei that underwent meiotic S-phase in the presence of SCC-2 plus a population of early prophase nuclei that lacked SCC-2 from the onset of meiosis. We first confirmed that 14 hours of auxin treatment induced efficient depletion of SCC-2::AID::GFP from all meiotic nuclei (Figure 2B). We then stained auxin-treated germlines with anti-COH-3/4 antibodies and observed that SCC-2 depletion induced a strong reduction (38%) of COH-3/4 signal in pachytene nuclei (Figure 2B), which underwent S-phase in the presence of SCC-2. In contrast, SCC-2 depletion resulted in a moderate decrease (9%) of REC-8 staining intensity in pachytene nuclei (Figure 2B). Importantly, SCC-2 depletion for 14 hours completely eliminated REC-8 staining from axial elements of early prophase nuclei (Figure 2C), consistent with the requirement of SCC-2 in the initial loading of REC-8 cohesin (Lightfoot et al., 2011), and reinforcing the idea that SCC-2-dependent loading of REC-8 cohesin in pachytene nuclei is minimal or not present. Thus, SCC-2 is required for maintaining the levels of chromosome-bound COH-3/4, but not REC-8, cohesin in pachytene nuclei.

Similar to the situation in *C. elegans*, mouse NIPBL (SCC2) localizes to axial elements of pachytene nuclei (Kuleszewicz et al., 2013), suggesting that cohesin turnover during pachytene, presumably of non-cohesive RAD21L complexes, may be a conserved feature of meiosis. This is further supported by observations in *Drosophila*, where the C(2)M kleisin, which does not provide SCC, is incorporated into axial elements during pachytene (Gyuricza et al., 2016). The localisation of SCC-2/NIPBL to pachytene axial elements may also indicate that active loop extrusion takes place at this stage, as Scc2 is required for activating cohesin’s ATPase activity and for loop extrusion in vitro (Davidson et al., 2019; Petela et al., 2018). Moreover, during the G1 phase of the mitotic cell cycle in yeast, Scc2 prevents removal of cohesin by a Wapl-independent mechanism (Srinivasan et al., 2019). Whether a similar mechanism may operate during meiotic prophase is not known, but our observations that SCC-2 localises to pachytene axial elements and that its depletion reduces axis-associated COH-3/4 cohesin would be consistent with SCC-2 acting to prevent cohesin release. Clarifying the roles of SCC-2 in controlling cohesin function during meiotic prophase is an important goal for future studies.

### SCC is provided by REC-8 complexes

We and others have proposed that during *C. elegans* meiosis SCC is exclusively provided by REC-8 cohesin, while COH-3/4 complexes associate with individual chromatids to regulate higher-order chromosome structure (Crawley et al., 2016; Woglar et al., 2020). However, this possibility is largely inferred from observations in mutants in which only REC-8 or COH-3/4 cohesin is present and therefore in which chromosome morphogenesis was partially impaired. Moreover, an involvement of COH-3/4 cohesin in SCC has also been proposed (Severson and Meyer, 2014), and in the absence of both WAPL-1 and proteins required for crossover formation COH-3/4 appear to mediate inter-sister attachments in diakinesis oocytes (Crawley et al., 2016). A direct assessment of the contribution of REC-8 and COH-3/4 cohesin to SCC under normal conditions requires the ability to specifically remove these complexes in a temporally-resolved manner after normal chromosome morphogenesis. For example, in mouse oocytes arrested at metaphase I, TEV-mediated removal of REC8 revealed that SCC is exclusively provided by REC8 cohesin, despite the presence of SCC1 cohesin (Tachibana-Konwalski et al., 2010). We have previously shown that versions of REC-8::GFP and COH-3::mCherry carrying TEV recognition motifs in the central region of these kleisins are fully functional and allow kleisin cleavage following TEV protease micro injection into the germline (Castellano-Pozo et al., 2020). We took advantage of these strains to assess REC-8 and COH-3 contribution to SCC at different stages of meiosis. In the case of COH-3, experiments were performed in a *coh-4* mutant background due to functional redundancy between COH-3 and COH-4 (Severson et al., 2009). We first evaluated the impact of removing REC-8 or COH-3 from diakinesis oocytes, which contain 6 bivalents as a result of SCC and the presence of inter-homologue crossover events (Figures 3A-B). TEV-mediated removal of REC-8::GFP caused partial disassembly of diakinesis bivalents, which in most cases appeared as four lobed-structures that remained weakly connected at the crossover site, with each lobe likely corresponding to one of the four chromatids present in a bivalent (Figure 3A). We called these structures “lobed bivalents”. In addition to lobed bivalents, TEV-mediated REC-8 removal resulted in 35% of oocytes displaying individual chromatids that were fully detached (Figure 3A). A similar situation was observed when REC-8 was removed from diakinesis oocytes using auxin-mediated depletion (Figure S1A). Importantly, visualisation of COH-3 and condensin II confirmed that these SMC complexes remained bound to diakinesis chromosomes following REC-8 removal, including on fully detached chromatids (Figure S1B-C). Therefore, the observed disruption in SCC is specifically caused by the loss of REC-8 and not by an indirect effect of REC-8 removal on other SMC complexes.

**Figure 3.**
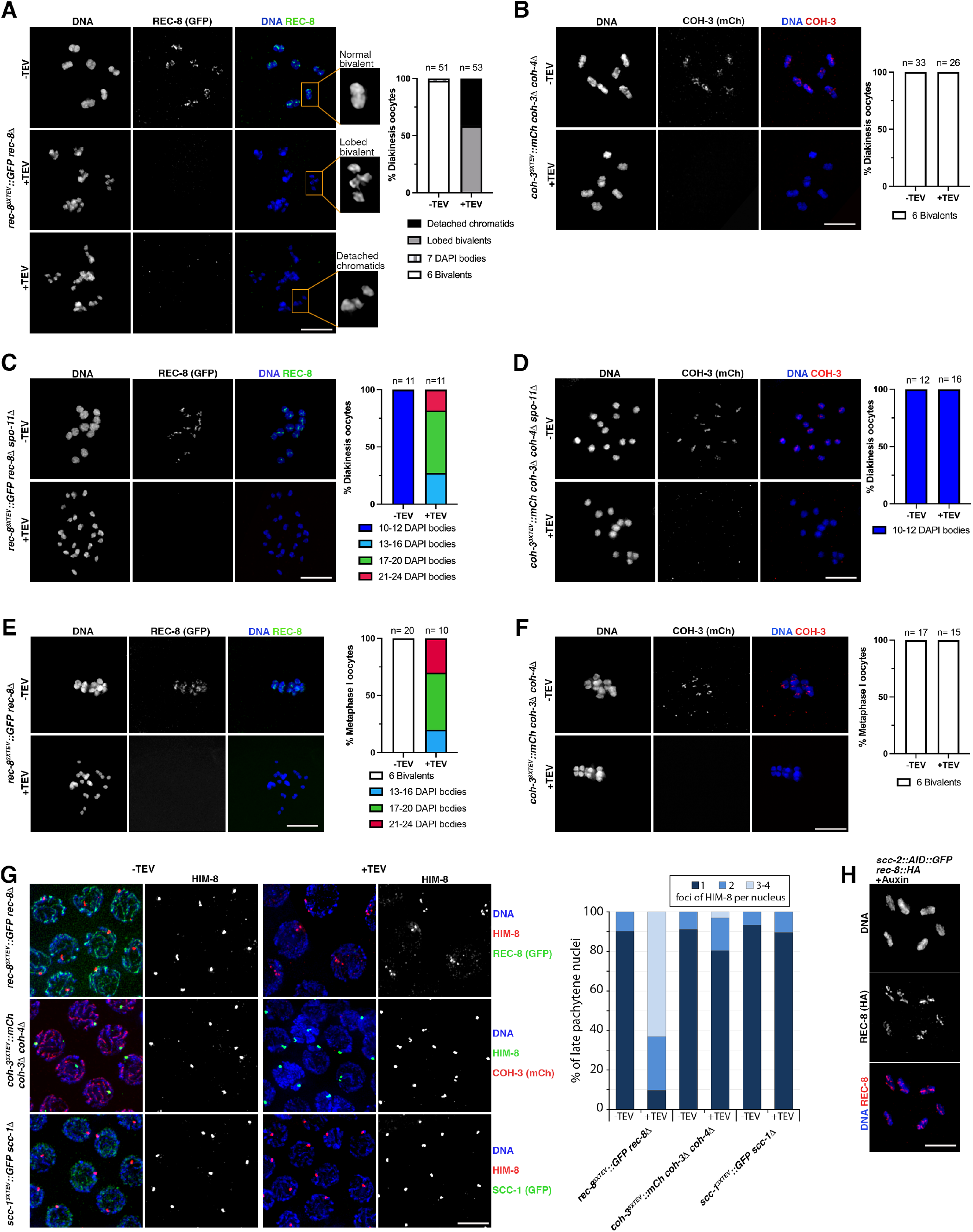
REC-8 complexes provide SCC. **(A-D)** Diakinesis oocytes of indicated genotypes stained with DAPI and anti-GFP (REC-8) or anti-mCherry (COH-3) antibodies from untreated controls (-TEV) and 3.5 hours post TEV-mediated kleisin removal. Note that REC-8 removal transforms bivalents into lobed structures (middle panel in (A)) and also induces appearance of detached chromatids (bottom panel in (A)). Bivalent morphology remains unaffected by COH-3 removal (B). See magnified bivalents in (A) for examples of the different morphological categories used for the quantification of diakinesis oocytes. (C-D) REC-8, but not COH-3, removal induces separation of sister chromatids in *spo-11* mutant oocytes. **(E-F)** Metaphase I-arrested oocytes (*apc-2* RNAi) stained with DAPI and anti-GFP (REC-8) or anti-mCherry (COH-3) antibodies from untreated controls (-TEV) and 3.5 hours post TEV-mediated kleisin removal. Note separation of sister chromatids following REC-8, but not COH-3, removal. **(G)** Late pachytene nuclei stained with DAPI anti HIM-8-antibodies and anti-GFP (REC-8 or SCC-1) or anti-RFP (COH-3) antibodies from untreated controls (-TEV) and 3.5 hours following TEV-mediated kleisin removal. Note that removal of REC-8, but not COH-3 or SCC-1, leads to the appearance of nuclei with 3 or 4 HIM-8 foci, indicating separation of sister chromatids. Number of nuclei analysed= REC-8 (92 -TEV, 92 +TEV), COH-3 (135 -TEV, 128 +TEV), SCC-1 (67 -TEV, 143 +TEV). **(H)** Diakinesis oocytes of *scc-2::AID::GFP rec-8::HA* worms exposed to auxin for 14 hours to induce SCC-2 depletion stained with DAPI and anti-HA (REC-8) antibodies. Note the presence of 6 bivalents displaying normal REC-8 staining and intact SCC. Scale bar= 5 *μ*m in all panels.

The fact that COH-3 remains associated with fully detached chromatids following REC-8 removal in diakinesis oocytes (Figure S1C) suggests that COH-3/4 complexes associate with individual chromatids and therefore do not partake in SCC. Indeed, TEV-mediated removal of COH-3 or the mitotic kleisin SCC-1, which also associates with meiotic prophase chromosomes (Severson and Meyer, 2014), caused no obvious morphological changes in diakinesis bivalents (Figures 3B and S1D), consistent with COH-3/4 or SCC-1 not providing SCC in diakinesis oocytes. We also attempted to simultaneously remove all cohesin variants by inducing auxin-mediated depletion of SMC-1, a subunit common to all cohesin complexes. SMC-1 depletion produced the appearance of lobed bivalents and detached chromatids in diakinesis oocytes (Figure S1E), as observed following REC-8 depletion. We noted that in most oocytes a small pool of SMC-1 signal remained associated with the crossover site (Figure S1E). We have previously observed that a population of REC-8 cohesin associated with crossover sites in late pachytene chromosomes is partially resistant to TEV-mediated removal (Castellano-Pozo et al., 2020). Thus, it is possible that a feature of chromosome structure present in the vicinity of crossover sites makes cohesin complexes associated with these regions more resistant to removal by the methods used here. To test if a recombination-dependent feature could explain the weak inter-chromatid connections present in the lobed diakinesis bivalents induced by REC-8 removal, we removed REC-8 from diakinesis oocytes of *spo-11* mutants, which lack crossovers due to the absence of DNA double strand breaks (DSBs) that initiate meiotic recombination (Dernburg et al., 1998). In this case, REC-8 removal induced a more penetrant loss of SCC (Figure 3C), suggesting that a recombination-dependent feature of chromosome organisation is involved in the weak chromatid connections observed when REC-8 is removed from diakinesis bivalents. We also tested a possible role of COH-3 in mediating SCC in diakinesis bivalents of *spo-11* mutants, but in this case, SCC remained intact following COH-3 removal (Figure 3D). These results suggest that SCC is exclusively provided by REC-8 complexes in diakinesis oocytes.

Next, we focused our attention to SCC in oocytes at the metaphase I stage, when SCC is required to ensure correct orientation of bivalents on the spindle. Removal of REC-8 from metaphase I–arrested oocytes caused a general loss of SCC, with most oocytes displaying between 17 and 24 DAPI-stained bodies (full loss of SCC would result in 24 chromatids), while 6 bivalents remained present following COH-3 or SCC-1 removal (Figures 3E-F and S1F). These results are consistent with REC-8 complexes providing SCC in metaphase I oocytes, and suggest that the weak chromatid attachments that remain in diakinesis bivalents following REC-8 removal are resolved in metaphase I oocytes. This may be due to the assembly of the first meiotic spindle, which may be sufficient to pull apart weakly attached chromatids, or to the resolution of a crossover-associated chromosome structure between diakinesis and metaphase I. Regardless, the clear conclusion from our temporally-resolved kleisin removal experiments is that, similar to mouse oocytes (Tachibana-Konwalski et al., 2010), SCC is exclusively provided by REC-8 cohesin in metaphase I oocytes of *C. elegans*.

Since SCC is established during meiotic S-phase and must persist until the meiotic divisions, and given that REC-8 provides SCC during metaphase I, we hypothesised that SCC is provided by REC-8 cohesin throughout meiotic prophase. We tested this possibility by monitoring SCC at the chromosomal end of the X chromosomes bound by the HM-8 protein (Phillips et al., 2005). Imaging of HIM-8 foci in late pachytene nuclei demonstrated that removal of REC-8, but not of COH-3 or SCC-1, caused loss of SCC (Figure 3G). These observations suggest that SCC is provided by REC-8 complexes throughout meiosis, consistent with a model in which complexes that provide SCC are stably bound to chromosomes between DNA replication and the onset of the meiotic divisions (Burkhardt et al., 2016). The stable association of cohesion-providing REC-8 complexes with meiotic chromosomes is also supported by our findings that SCC-2 is not required for maintaining REC-8 on pachytene chromosomes. Moreover, we found that both REC-8 and SCC remained intact in diakinesis oocytes after 14 hours of auxin-mediated SCC-2 depletion (Figure 3H). In contrast, maintenance of COH-3/4 complexes, presumably associated with individual chromatids, does require SCC-2 in pachytene nuclei (see Figure 2B). These differences between REC-8 and COH-3 in terms of their participation in SCC and their dependency on SCC-2 to sustain chromosomal association are reminiscent of observations in yeast mitotic cells, where Scc2 is essential for loading cohesin and for maintaining cohesin’s association with unreplicated DNA, but has no role in maintaining cohesion during G2/M phases (Ciosk et al., 2000; Srinivasan et al., 2019).

### REC-8 cohesin plays a more prominent role in DNA repair than COH-3/4 cohesin

Cohesin is thought to have diverse functions in the repair of DSBs, including a SCC-dependent role in homologous recombination (Sjogren and Nasmyth, 2001), and roles in regulating homology search of DSB ends and in the formation of repair foci, which are proposed to depend on the loop extrusion activity of cohesin (Arnould et al., 2021; Piazza et al., 2021). Our results so far show that stably-bound REC-8 cohesin provides SCC while a larger population of dynamically associated COH-3/4 cohesin orchestrates chromosome organisation. We sought to clarify whether these differential activities correlate with the ability of REC-8 and COH-3/4 cohesin to promote DSB repair. Analysis of chromosome morphology in diakinesis oocytes after exogenous DSBs are introduced by ionizing radiation (IR) provides a clear read out of DSB repair capability, as impaired DSB repair during pachytene leads to chromosome fragmentation in diakinesis oocytes. Chromosome fragments were largely lacking from diakinesis oocytes of wild-type worms exposed to 100 Gy of IR, evidencing efficient DSB repair (Figure 4A). In contrast, extensive chromosome fragmentation was evident in diakinesis oocytes of irradiated r*ec-8* mutants, with around 40% of the DAPI-stained bodies displaying an area consistent with chromosome fragments (Figure 4A). Chromosome fragmentation was also detected in diakinesis oocytes of irradiated *coh-3 coh-4* double mutants, although to a much lesser extent than in *rec-8* mutants. We also evaluated IR-induced DSB repair in pachytene nuclei by visualising recombination intermediates containing RAD-51 in backgrounds lacking SPO-11 and therefore endogenous DSBs. 24 hours after exposure to 10Gy, which induces a large accumulation of RAD-51 foci (Lightfoot et al., 2011), low numbers of RAD-51 foci were present in late pachytene nuclei of *spo-11* and *spo-11 coh-3 coh-4* mutants, consistent with efficient DSB repair (Figure 4B). In contrast, late pachytene nuclei of irradiated *rec-8* mutants contained high numbers of RAD-51 foci, evidencing impaired DSB repair (Figure 4B). Thus, SCC-providing REC-8 complexes play a more prominent role in DSB repair than non-cohesive, dynamically bound, COH-3/4 complexes. This is despite the fact that COH-3/4 complexes are much more abundant than REC-8 complexes and that they are essential for the integrity of axial elements (Figure 1B). Moreover, in pachytene nuclei of *rec-8* mutants, most sister chromatids are paired up due to the assembly of inter-sister synaptonemal complex (Cahoon et al., 2019), a phenomenon also observed in mouse *Rec8* mutants (Xu et al., 2005). However, our results demonstrate that inter-sister connections mediated by the synaptonemal complex in *rec-8* mutants are not capable of supporting efficient repair of IR-induced DSBs, suggesting that the function of REC-8 cohesin in DSB repair, at least for IR-induced DSBs, is mechanistically coupled to its role in SCC.

**Figure 4.**
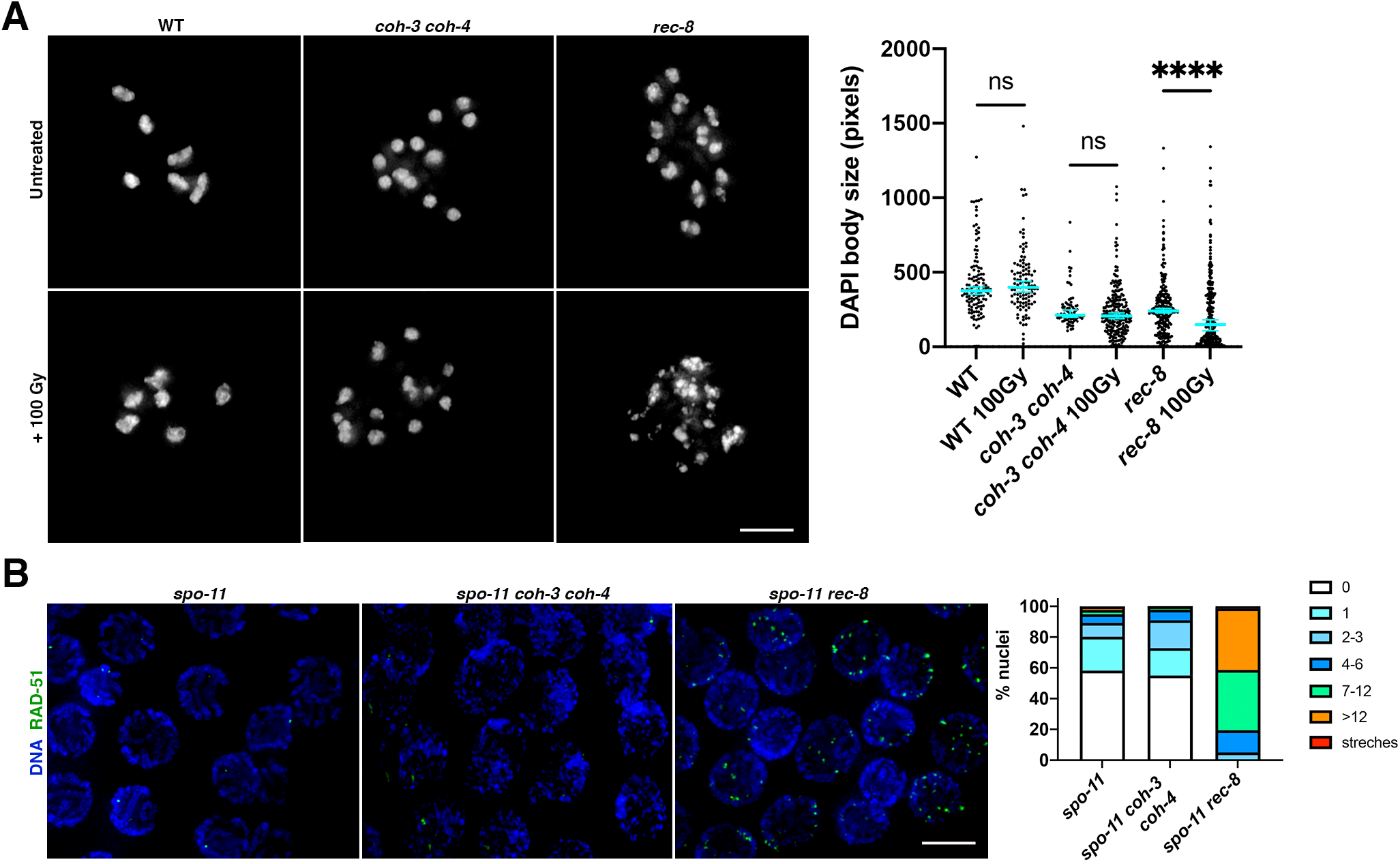
REC-8 complexes promote DSB repair. **(A)** Diakinesis oocytes of indicated genotypes stained with DAPI in untreated controls and 26 hours after worms were exposed to 100 Gy of IR. Note the extensive appearance of small chromatin bodies in *rec-8* mutants, which indicate chromosome fragmentation. Graphs show the distribution of DAPI-stained bodies with a given area in full nucleus projections of diakinesis oocytes. Number of oocytes: WT n=22 oocytes (0 Gy) n=19 (100Gy); *rec-8* n=20 (0 Gy) n=16 (100 Gy); *coh-3 coh-4* n=7 (0 Gy) n=22 (100Gy). Error bars indicate median with 95% CI, p values were calculated by two-tailed Mann-Whitney test. **(B)** Late pachytene nuclei of indicated genotypes stained with DAPI and anti-RAD-51 antibodies 24 hours after worms were exposed to 10 Gy of IR. Note high number of RAD-51 foci in *rec-8 spo-11*, but not in *spo-11 coh-3 coh-4* or *spo-11* mutants. Number of nuclei analysed= 298 (*spo-11*), 276 (*spo-11 coh-3 coh-4*), 237 (*spo-11 rec-8*). Scale bar= 5 *μ*m in all panels.

## Conclusions

We reveal a clear distribution of functions between stably-bound and low abundance REC-8 complexes, which provide SCC and DSB repair, and high-abundance COH-3/4 complexes that ensure the integrity of axial elements and associate dynamically with pachytene chromosomes in a process controlled by WAPL-1 and SCC-2. Our studies support functional conservation between worm and mouse REC8 cohesin, with REC8 complexes providing SCC and being largely refractory to WAPL-mediated removal in both organisms (Crawley et al., 2016; Silva et al., 2020; Tachibana-Konwalski et al., 2010), and suggest that functional conservation also likely extends to COH-3/4-RAD21L complexes. Therefore, we hypothesise that RAD21L complexes associate dynamically with meiotic chromosomes regulated by WAPL and NIPBL. We support a model in which REC-8 complexes loaded during meiotic S-phase tether sister chromatids until the meiotic divisions, while COH-3/4 are loaded post S-phase on individual chromatids to control higher-order chromosome structure, likely by performing loop extrusion. Mitotic yeast and human cohesin complexes containing the Scc1 kleisin mediate both SCC and loop extrusion, but these processes are thought to involve different modes of DNA-cohesin interactions and are therefore proposed to be mutually exclusive (Davidson et al., 2019; Srinivasan et al., 2019). How different populations of Scc1 cohesin are regulated to perform either SCC or loop extrusion is not understood, but our findings suggest that during meiosis in higher eukaryotes these activities are determined by kleisin identity.

## Methods

### *C. elegans* genetics and growing conditions

All strains were maintained on *E. coli* (OP50) seeded NG agar plates at 20 °C under standard conditions. N2 Bristol strain was used as the wild-type strain. All studies were performed using young adults at 18-24 hours post L4-larvae stage unless indicated. The following mutant alleles were used: *wapl-1(tm1814), rec-8(ok978), coh-3(gk112), coh-4(tm1857), scc-1(ok1017)*, and *spo-11(ok79)*. Table S1 contains a full list of the strains used in this study.

### Generation of transgenic *C. elegans* strains

For the generation of transgenic strains carrying single copy insertions of the desired transgene, we used a strain carrying the *ttTi5605* MosSCI transposon insertion on chromosome II, except for transgene *fqSi18*, which was inserted at the *ttTi4348* locus on chromosome I. Transgene insertion was performed using the protocol described in (Frokjaer-Jensen et al., 2008). The *scc-1*^*3XTEV*^*::GFP* transgene was generated by adding a 75 bp fragment encoding for 3 repeats of the TEV recognition motif (ENLYFQGASENLYFQGELENLYFQG) after SCC-1’s E293 codon in a vector expressing SCC-1::GFP under the *scc-1* promoter and 3’ UTR. Table S2 contains a list of transgenes used in this study.

CRISPR-mediated genome editing was performed as described in (Paix et al., 2017), using preassembled Cas9-sgRNA complexes, single-stranded DNA oligos as repair templates, and *dpy-10* as a co-injection marker. To generate the *scc-2::AID::GFP* allele we introduced 105 bp encoding the 35 amino acids of the AID tag (Zhang et al., 2015) between the last codon of *scc-2* and the start codon of GFP. The *rec-8::HA* allele was generated by introducing an 81 bp fragment encoding for three repeats of the HA tag (YPYDVPDYAYPYDVPDYAYPYDVPDYA) before *rec-8*’s STOP codon, while the *rec-8*^*3XTEV*^ alle was generated by introducing a 75 bp fragment encoding for 3 repeats of the TEV recognition motif (ENLYFQGASENLYFQGELENLYFQG) after *rec-8*’s Q289 codon. The *rec-8::GFP, coh-3::GFP*, and *coh-4::GFP* alleles were generated by SunyBiotech and all contained a C-terminal GFP sequence containing three artificial introns.

### Auxin-mediated protein degradation

All strains used for auxin-mediated protein degradation were homozygous for the *ieSi38* transgene (Table S2) expressing the TIR1 protein under the *sun-1* promoter and all proteins targeted for auxin-mediated degradation were expressed fused to the 35 amino acid AID tag (Zhang et al., 2015). Auxin treatment was performed by placing young adult worms, at 18-24 hours post L4-larvae stage, in seeded NG agar plates containing 4 mM Auxin for the indicated periods of time.

### TEV protease microinjection

Germline injections were performed as described in (Castellano-Pozo et al., 2020) using a Narishige IM-31 pneumatic microinjector attached to an inverted Olympus IX71 microscope. Needles were made using a micropipette puller P-97 (Intracell) and borosilicate glass filaments with a 1.0 mm O.D. and 0.58 mm I.D. (BF100-58-10, Sutter Instruments). AcTEV™ Protease (Thermo Fisher, Cat. No. 12575) was used in a mix containing 10U/μl TEV protease in 50 mM Tris-HCl, pH 7.5, 1 mM EDTA, 5 mM DTT, 50% (v/v) glycerol, 0.1% (w/v) Triton X-100.

### Immunostaining and image acquisition

Germ lines from young adult hermaphrodites were dissected, fixed and processed for immunostaining as described in (Castellano-Pozo et al., 2020). Briefly, worms were dissected in EGG buffer (118 mM NaCl, 48 mM KCl_2_, 2mM CaCl_2_, 2mM MgCl_2_, 5mM HEPES) containing 0.1% Tween and fixed in the same buffer containing 1% paraformaldehyde for 5 minutes. Slides were immersed in liquid nitrogen before removing the coverslip and then placed in methanol at -20 °C for 5 minutes, followed by 3 washes of 10 minutes each in PBST (1x PBS, 0.1% Tween) and blocking in PBST 0.5% BSA for 1 hour. Following incubation over night with primary antibodies (see Supplemental methods) diluted in PBST, slides were washed three times for 10 minutes each in PBST. Slides were then incubated in the dark at room temperature for 2 hours with secondary antibodies diluted in PBST, washed with PBST three times for 10 minutes each and counterstained with DAPI. Finally, slides were washed for 1 hour in PBST and mounted using Vectashield (Vector). Images were acquired as 3D stacks using a 100X lens in a Delta Vision Deconvolution system equipped with an Olympus 1×70 microscope and images were deconvolved using SoftWoRx 3.0 (Applied Precision) and mounted using Photoshop.

### Super resolution structured illumination microscopy

For images shown on Figure 1C, immunostaining was performed as described above, but slides were mounted using ProLong Diamond mounting media and covered with with Zeiss high-performance 0.17± 0,005 coverslips. Images were acquired using a Zeiss Elyra S1 microscope, processed with Fiji, and mounted in Photoshop.

## See supplemental Methods for

**Quantification of microscopy images, Gamma irradiation, RNA interference, and FRAP protocols**.

## Acknowledgements

We thank D. Dormann and C. Whilding from the MRC LMS microscopy facility for help with analysis of FRAP, Nicola Silva for providing anti-RAD-51 antibodies, and Barbara Meyer for providing anti-HCP-6 antibodies. This work was supported by a MRC core-funded grant to E.M.-P. (MC-A652-5PY60), postdoctoral Fellowships from Fundación Alfonso Martin Escudero and EMBO to M.C.-P, a grant from Universidad Autónoma de Madrid to J.P. and P.L-J. (BIOUAM02-2020), and an EMBO scientific exchange grant to P.L.-J.

