## Supplemental Information for "The kleisin subunit controls the function of meiotic cohesins by determining the mode of DNA binding and differential regulation by SCC-2 and WAPL-1"

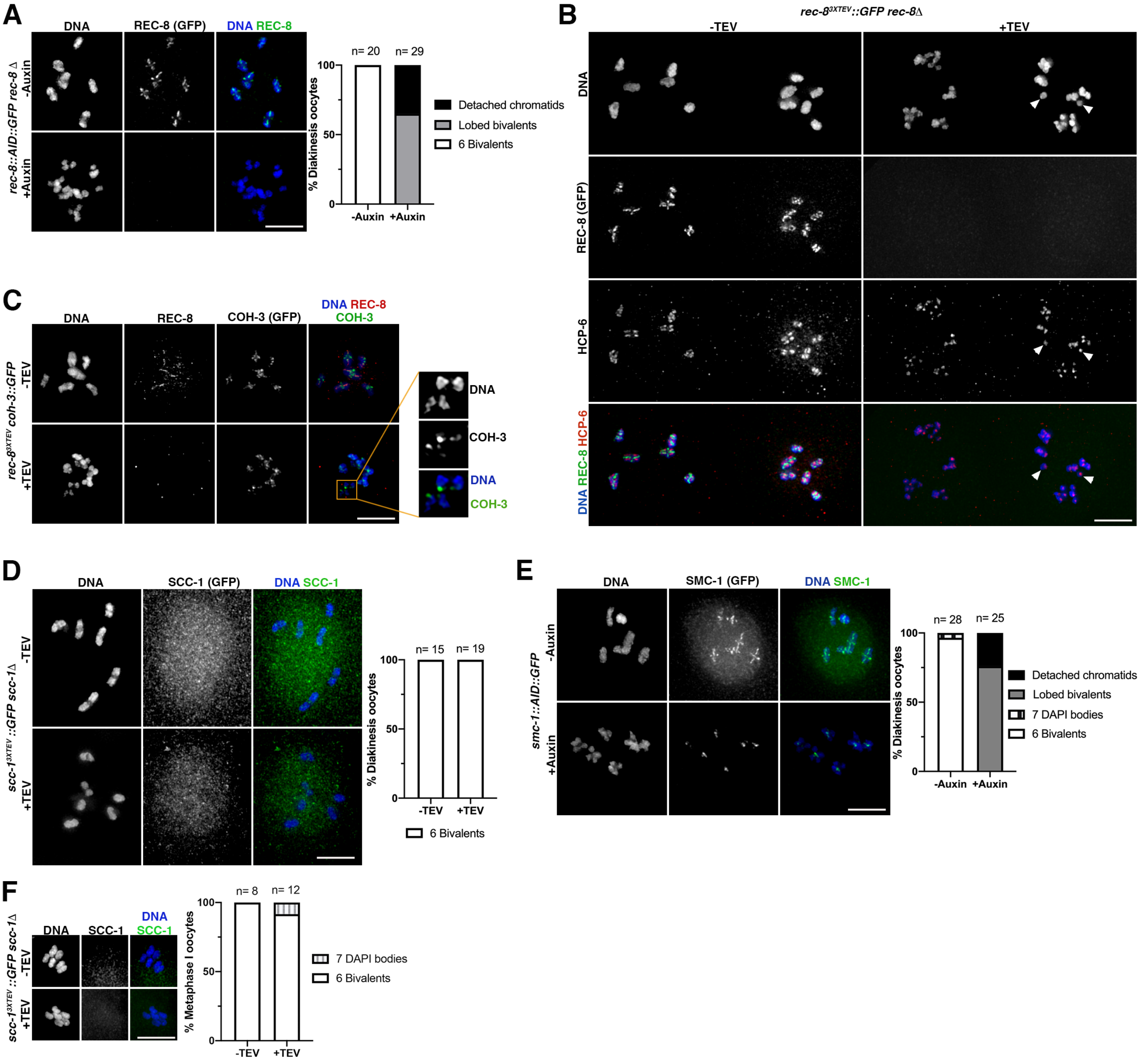


**Figure S1. (A)** Diakinesis oocytes of *rec-8::AID::GFP* worms stained with DAPI and anti-GFP antibodies from control worms (-Auxin) or from worms exposed to auxin for 8 hours (+Auxin), which induces loss of GFP signal (REC-8) and the appearance of lobed bivalents and detached chromatids. **(B)** Diakinesis oocytes of *rec-8$^{3XTEV}$::GFP rec-8Δ* worms stained with DAPI, anti-GFP and anti-HCP-6 (condensin II) antibodies from untreated controls (-TEV) and 3.5 hours after TEV-mediated REC-8 removal. Note that HCP-6 signals remain associated with both partially and fully separated (arrowheads) chromatids following REC-8 removal. **(C)** Diakinesis oocytes of *rec-8$^{3XTEV}$ coh-3::GFP* worms (both alleles generated by CRISPR) stained with DAPI, anti-GFP and anti-REC-8 antibodies from untreated controls (-TEV) and 3.5 hours after TEV-mediated REC-8 removal. Note that COH-3 signal remains associated with chromosomes following REC-8 removal, including with fully detached chromatids (inset). **(D)** Diakinesis oocytes of *scc-1*$^{3XTEV}$*::GFP scc-1Δ* worms stained with DAPI, anti-GFP antibodies from untreated controls (-TEV) and 3.5 hours after TEV-mediated SCC-1 removal. Bivalents remain intact in TEV-treated oocytes. **(E)** Diakinesis oocytes of *smc-1::AID::GFP* worms stained with DAPI and anti-GFP antibodies from control worms (-Auxin) or from worms exposed to auxin for 8 hours. Note that auxin treatment results in the appearance of lobed bivalents and large loss of GFP signal, apart from small signals associated with the chiasma site. **(F)** Metaphase I-arrested oocytes of *scc-1*$^{3XTEV}$*::GFP scc-1Δ* worms stained with DAPI and anti-GFP antibodies from untreated controls (-TEV) and 3.5 hours after TEV-mediated SCC-1 removal. Bivalents remain intact following SCC-1 removal. Scale bar= 5 μm in all panels.

| Strain | Genotype | Origin |
|---|---|---|
| ATG473 | *rec-8(syb803 [rec-8::GFP]) IV* | This study |
| ATG472 | *coh-3(syb751 [coh-3::GFP]) V* | This study |
| ATG556 | *coh-4(syb1273 [coh-4::GFP]) V* | This study |
| ATG228 | *smc-1 (fq20 [smc-1::GFP] I* | Crawley |
| ATG252 | *smc-1 (fq20 [smc-1::GFP] I; rec-8(ok978) IV / nT1 [qls51] (IV;V)* | This study |
| ATG253 | *smc-1 (fq20 [smc-1::GFP] I; coh-4(tm1857) coh-3(gk112) V / nT1 [qls51] (IV;V)* | This study |
| ATGSi23 | *fqSi23 II; rec-8 (ok978) IV* | Crawley et al 2016[1] |
| ATGSi191 | *fqSi18 I; fqSi23 II; rec-8 (ok978) IV; coh-3 (gk112) V* | This study |
| ATG571 | *wapl-1(tm1814) rec-8(syb803 [rec-8::GFP]) IV /nT1 [unc-? (n754) let-? qls50] (IV;V)* | This study |
| ATG570 | *wapl-1(tm1814) IV; coh-3(syb751 [coh-3::GFP]) V /nT1 [unc-? (n754) let-? qls50] (IV;V)* | This study |
| ATG572 | *wapl-1(tm1814) IV; coh-4(syb1273 [coh-4::GFP]) coh-3(syb751 [coh-3::GFP]) V //nT1 [unc-? (n754) let-? qls50] (IV;V)* | This study |
| ATG282 | *scc-2(fq23 [scc-2::AID::GFP]) II ; ieSi38 IV* | This study |
| ATG693 | *scc-2(fq23 [scc-2::AID::GFP]) II ; ieSi38 rec-8(fq169[rec-8::3XHA]) IV* | This study |
| ATGSi355 | *fqSi16 II; rec-8(ok978) IV* | Castellano-Pozo et al 2020[2] |
| ATGSi441 | *fqSi15 II ; coh-4(1857) coh-3(gk112) V* | Castellano-Pozo et al 2020[2] |
| ATGSi392 | *fqSi16 II ; rec-8(ok978) spo-11(ok79) IV / nT1 [qls51] (IV;V)* | This study |
| ATGSi470 | *fqSi15 II; spo-11(ok79) IV; coh-4(1857) coh-3(gk112) V / nT1 [unc-? (n754) let-? qls50] (IV;V)* | This study |
| ATG323 | *fqSi17 II ; ieSi38 rec-8(ok978) IV* | Castellano-Pozo et al 2020[2] |
| ATG541 | *rec-8(fq32[rec-8::TEV] IV; coh-3(syb751[coh-3::GFP])* | This study |
| ATG415 | *smc-1 (fq64[smc-1::AID::GFP]) I; ieSi38 IV* | Castellano-Pozo et al 2020[2] |
| TY5120 | *coh-4(tm1857) coh-3(gk112) V/nT1 [qls51] (IV;V)* | CGC |
| VC666 | *rec-8(ok978) IV/nT1 [qls51] (IV;V)* | CGC |
| AV106 | *spo-11(ok79) IV/nT1 [unc-? (n754) let-?] (IV;V)* | CGC |
| ATG137 | *rec-8(ok978) IV; spo-11(ok79) IV/ nT1 [qls51] (IV;V)* | Crawley et al 2016[1] |
| ATG213 | *spo-11(ok79) IV; coh-4(tm1857) coh-3(gk112) V/ nT1 [unc-? (n754) let-?] (IV;V)* | Crawley et al 2016[1] |

**Table S1. *C. elegans* strains used in this study**

| Transgene | Genotype |
|---|---|
| *fqSi23* | [*Prec-8 rec-8::GFP 3'UTR rec-8*; *cb-unc-119(+)*] |
| *fqSi18* | [*Pcoh-3 coh-3::mCherry 3'UTR coh-3*; *cb-unc-119(+)*] |
| *ieSi38* | [*Psun-1 TIR1::mRuby 3'UTR sun-1; cb-unc-119(+)*] |
| *fqSi16* | [*Prec-8 rec-8*$^{3XTEV}$*::GFP 3'UTR rec-8*; *cb-unc-119(+)*] |
| *fqSi15* | [*Pcoh-3 coh-3::*$^{3XTEV}$*::mCherry 3'UTR coh-3*; *cb-unc-119(+)*] |
| *fqSi17* | [*Prec-8 rec-8::AID::GFP 3'UTR rec-8*; *cb-unc-119(+)*] |

**Table S2. Transgenes used in this study**

### Supplemental Methods

#### Primary antibodies used

The following primary antibodies and dilutions were used: goat anti-GFP-488-conjugated (1:200) (Roche), rat anti-mCherry (1:1000) (5F8, Chromotek), rabbit anti-COH-3/4 (1:400) [1], mouse anti-REC-8 (1:100) (Novus Biologicals), guinea pig anti-HCP-6 (1:400) [3], rabbit anti-HIM-8 (1:500) (Novus Biologicals), and rabbit anti-RAD-51 [4].

#### Scoring number of DAPI-stained bodies in diakinesis and metaphase I oocytes

Worms of indicated genotypes and treatments were dissected and processed for immunostaining as described in main methods, including the final DAPI staining step. Images were acquired as 3D stacks using a 100X lens in a Delta Vision Deconvolution system equipped with an Olympus 1X70 microscope. Images were deconvolved using SoftWoRx 3.0 (Applied Precision) and mounted in Photoshop. The number and appearance of DAPI-stained bodies in diakinesis and metaphase I oocytes was scored in 2D projections of three-dimensional intact germlines/embryos. Projections of diakinesis and metaphase I oocytes result in some overlap of DAPI-stained bodies, especially in situations where loss of SCC results in high numbers of DAPI-stained bodies, therefore in some cases the number of DAPI-stained bodies my represent a slight underestimation of the overall number of individual chromatin bodies present.

The area in pixels of DAPI-stained bodies in diakinesis oocytes (Figure 4A) was calculated by generating maximum intensity projections of diakinesis nuclei and using CellProfiler to obtain the size in pixels of individual DAPI-stained bodies identified in 2D projections.

#### Gamma irradiation

Worms were irradiated in a IBL 637 cell irradiator containing a caesium-137 source. NGM plates, containing young adult worms, were directly irradiated for the appropriate amount of time, resulting in irradiation of 100Gy or 10Gy, as required.

**Metaphase I arrest by RNA interference (RNAi) of *apc-2***

Metaphase I arrest was achieved by downregulation of the anaphase promoting complex (APC) via RNAi feeding using the *apc-2* clone from the Ahringer library (HT115 bacteria transformed with a vector for IPTG-inducible expression of dsRNA). Bacteria containing the *apc-2* vector, as well as empty vector (HT115) control, were both grown overnight at 37 °C in LB with 50 µg/ml ampicillin. Cultures were collected and seeded onto NGM agar plates containing 1 mM IPTG and 25 µg/ml ampicillin. Plates were incubated overnight at 37 °C to induce the expression of dsRNA. 18-24 hours post-L4 worms were then placed and experiments were performed after 48 hours.

**Quantitative analysis of fluorescence intensity**

To compare the occupancy of GFP-tagged meiotic kleisin subunits (Figure 1B), the peak axis fluorescence was measured. Dissected gonads of strains carrying GFP-tagged versions (generated by CRISPR) of REC-8, COH-3 and COH-4 were stained with FITC-conjugated aGFP antibodies. Acquisition was carried out on the Deltavision microscope, with a set exposure time. Sum-projections of the raw images were made on ImageJ to only include the half top or bottom of each nucleus (6-slice projections). Every nucleus was examined separately. To measure the peak axis intensity, a line was drawn over at least 2 clear axes of the nuclei, and the line profile was generated using the built-in "Plot profile" function of ImageJ. Peak values were called using an online-available macro (found here: https://www.imperial.ac.uk/medicine/facility-for-imaging-by-light-microscopy/equipment/software---fiji/). The macro used was the "Intensity" macro (Maxima and Minima of line profile Tool). Fluorescence measurements (in arbitrary units) were collected and the raw values were used to compare the different tagged protein occupancy by calculating the relative ratio of the proteins on the meiotic axis.

We used whole nucleus fluorescence quantification to compare the immunostaining intensity of REC-8::HA and COH-3/4 in untreated controls and worms in which SCC-2::AID::GFP was depleted by auxin treatment. Images were acquired on a Delta Vision microscope as 3D stacks using the same exposure settings in auxin-treated and untreated controls. For comparing fluorescence levels, non-deconvolved images were analysed in imageJ. Nuclei of interest were manually circled using the "oval" tool, one nucleus at a time, and the fluorescence of each slice was measured. The mean fluorescence of that nucleus was then calculated, after normalizing for the number of z-stack slices and the area of the circle drawn. Normalized, mean fluorescence values were directly compared between control and mutant strains, as arbitrary units.

**Fluorescence recovery after photobleaching (FRAP)**

The FRAP method used here was modified from Nadarajan et al., 2017 [5] and Pattabiraman et al., 2017 [6]. Young adult worms were used for all FRAP experiments and immobilized on 5% agarose pads for imaging, covered in a solution of 2mM levamisole in M9 medium. Imaging was performed in a Leica TCS SP5 system using the FRAP wizard (LASAF software), which allows for photobleaching of the areas of interest (ROIs). The FRAP wizard was only used for the pre-bleach, photobleaching and the 0min post-bleach images. Images were taken in 2D every minute using the 60X oil immersion lens, as 3D stack images were found to increase photobleaching in this setup. Nuclei of mid-pachytene were selected that at least 2 axes were visible at the top of the gonad, where resolution was better. ROIs for photobleaching were designed to span at least 2 axes, with the ROI size being kept similar throughout experiments. 1 or 2 nuclei were bleached for every gonad, but only one nucleus per gonad was analysed for curve fitting. For acquisition, the following settings were used: 100Hz scanning speed, 2 times line averaging, 1AU pinhole. A 488 Argon laser was used at 30% power and 15% sub-power, and a HyD filter was used, set at 502-552nm range. Bleaching was performed with the same laser at 60% sub-power, for 100msec. Images were exported at .lif files, and were analysed using ImageJ (version 2.0.0-rc-59/1.51k). The ImageJ-built in StackRreg algorithm was used to align the frames of the time-series.

Aligned time-series images were analysed as described in Nadarajan et al 2017 [5] and Pattabiraman et al., 2017 [6]. In short, 3 ROIs of interest were designed: Bleached area of the axis (ROI1), the whole nucleus (ROI2) and a background control, defined as a region in the gonad, but far enough from the bleached area (ROI3). Intensity measurements were carried out for all ROIs using ImageJ and data was analysed using Microsoft Excel. The analysis involved background subtraction (ROI3) from ROI1 and ROI2. The subtracted values were then normalised against each other, so that fluorescence loss in ROI2 was accounted for in ROI1 (adapted from Phair et al., 2004 [7]). The ROI1 post-bleach values were also normalised as relative intensities of the pre-bleach ROI1 values. The resulting “double-normalised” values were finally normalised again, this time by setting the initial post-bleach value to 1, by dividing all values by the initial difference [8].

Curve fitting was performed in GraphPad Prism software. Statistical analysis was performed using the one-phase association predicted $Y_{max}$ values (where the fitted curve is expected to plateau) and comparing them between different genotypes using the Mann-Whitney statistical test.

High resolution FRAP images (Figures 1E-F) were acquired on a Leica TCS SP5 system with the following settings: 100 Hz scanning speed, 3 times line averaging, 1 AU pinhole. REC-8::GFP was imaged with 488 laser at 30% overall power and 10% sub-power for acquiring, HyD detector set to 502 nm- 552 nm range at 300 gain. COH-3::mCherry was imaged with a 561 laser and 10% imaging power, HyD detector set to 590- 680 nm range, at 350 gain. Some manual adjustment of Z and X-Y focus was needed due to nuclear movement over long time points, this was done using very low intensity imaging (1400 Hz) to minimise acquisition bleaching. Images were exported as TIFFs.
